# Efficient mapping of the thalamocortical monosynaptic connectivity *in vivo* by tangential insertions of high-density electrodes in cortex

**DOI:** 10.1101/2022.12.28.522000

**Authors:** Jérémie Sibille, Carolin Gehr, Jens Kremkow

**Affiliations:** Neuroscience Research Center, Charité-Universitätsmedizin Berlin,10117 Berlin, Germany; Bernstein Center for Computational Neuroscience Berlin, 10115 Berlin, Germany; Institute for Theoretical Biology, Humboldt-Universität zu Berlin, 10115 Berlin, Germany; Einstein Center for Neurosciences Berlin, 10117 Berlin, Germany

## Abstract

Thalamus provides the principal input to cortex, thus understanding the mechanisms of cortical computations in behaving animals requires to characterize the thalamocortical connectivity *in vivo*. We show that tangential insertions of high-density electrodes into mouse cortex capture the activity of thalamocortical axons simultaneously with their synaptically connected cortical neurons. Multiple thalamic synaptic inputs to cortical neurons can be measured providing an efficient approach for mapping the thalamocortical connectivity *in vivo*.

## MAIN

The thalamus provides the main afferent input to all cortical regions and is crucial for its computations^1– 6^ and its functional organization^2,7^. However, how spikes from thalamic neurons are integrated into the ongoing cortical activity in behaving animals remains largely unknown due to technical difficulties in recording synaptically connected thalamocortical neuron pairs *in vivo*^8^. Electrophysiological paired recordings are the gold standard for characterizing monosynaptic connectivity *in vivo*^1,3,6,9^. This technique has revealed important insights into the organizing principles of the thalamocortical pathways^1,3–6^. However, *in vivo* paired recordings require a precise alignment between the electrodes in the thalamus and cortex to capture the activity of synaptically connected thalamocortical neuron pairs. This step is technically challenging and therefore limiting the numbers of identified connected neuron pairs per experiment^8^. Consequently, while important progress has been made over the last decades^1– 6,8,10^, our overall understanding of the thalamocortical processing in sensory and non-sensory areas remains incomplete and is an active field of research^11^.

A method capable of reliably recording the activity of synaptically connected thalamic and cortical neurons *in vivo* would greatly advance our understanding of thalamocortical processing and could reveal the mechanisms underlying perception and behavior. Recording the activity of thalamic neurons at the axonal level within cortex, simultaneously with the activity of nearby cortical neurons, would overcome the challenging step of aligning multiple probes across distant brain regions and allow to map the thalamocortical connectivity in a more efficient manner. Recently, we showed that high-density electrodes (Neuropixels probes^12^) capture the activity of retinal ganglion cell axons within the mouse superior colliculus, allowing a detailed characterization of the retinocollicular synaptic connectivity *in vivo*^13^. Here we investigated whether high-density electrodes can also record the activity of thalamic axons within cortex and thus allow to efficiently monitor and functionally map thalamocortical monosynaptic connections.

We now report that tangential insertions of Neuropixels probes into layer 4 of mouse visual cortex (V1) (Fig. 1g) can detect extracellular signals of thalamocortical axons (TCA)^14^ simultaneously with somatic action potentials of V1 neurons (V1N, Fig. 1a-c, cf. methods and Extended Data Fig. 1 for optimizing probe placement). To characterize the spatiotemporal properties of the TCA waveforms, we distinguished the axonal pre-synaptic contact field (AF) from its dendritic synaptic contact field (DF), quantifying their amplitudes and spatial spreads on the probe. Both AF and DF have smaller amplitudes compared to the waveforms of V1N (Fig. 1e, positive peak amplitude: V1N = 24.6 ± 16 µV, AF = 12.8 ±

**Figure 1:**
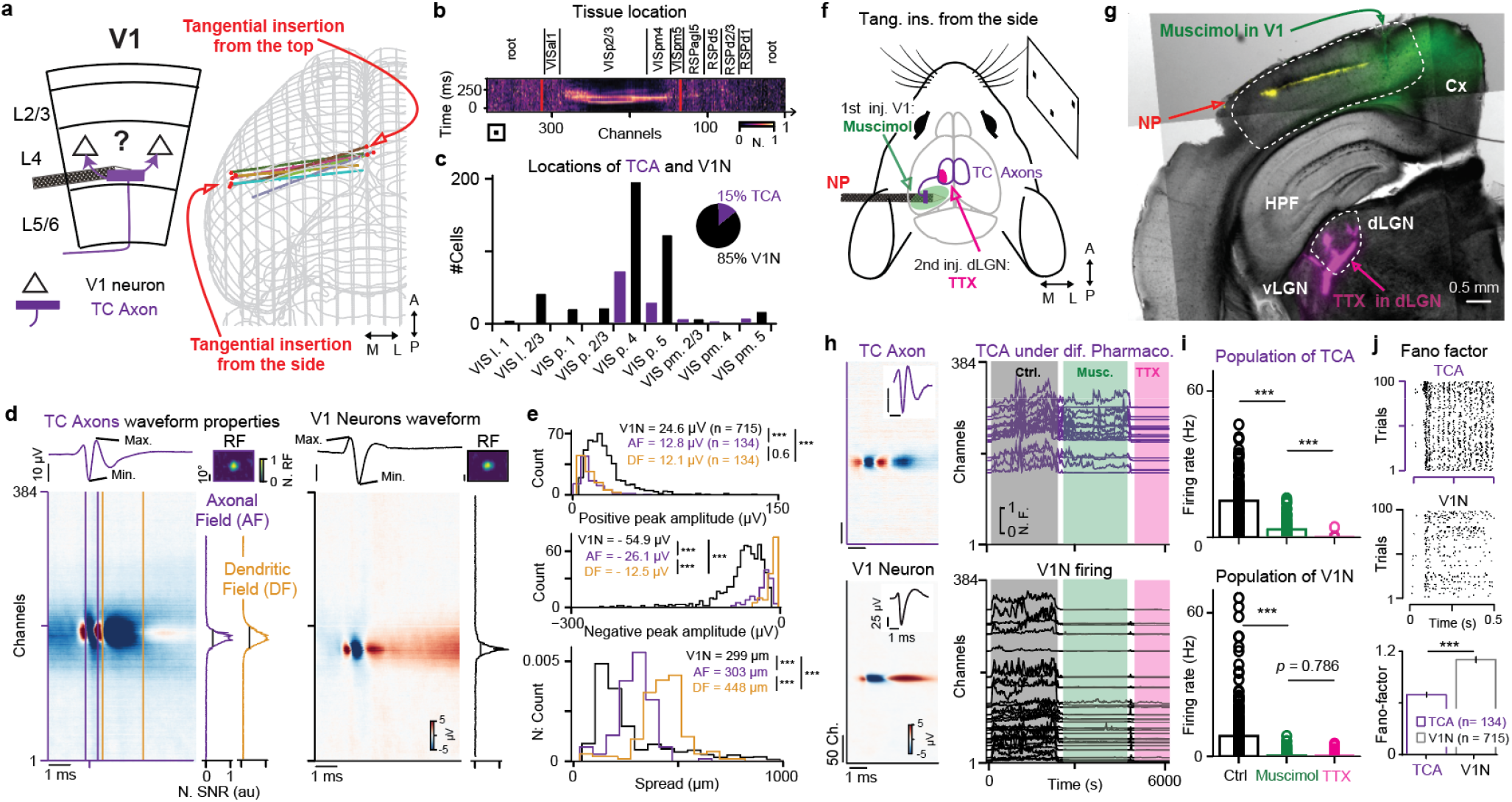
Neuropixels probes capture thalamocortical axons within cortex *in vivo*. **a**, Schematic of thalamocortical afferent (TCA) connecting cortical V1 neurons (V1N), and 3D rendering of half a mouse brain with the different probe tracks reconstructed for both the side and top insertions (right), *n =* 8 insertions, *n =* 8 mice. **b**, Reconstruction of one Neuropixels insertion via SHARP-track (top) aligned to the corresponding multi-unit activity (MUA, bottom). **c**, Quantification of the locations of TCA and V1N within cortical layers. **d**, Multi-channel waveform (MCW) of a TCA (left) and a V1N (right); TCA properties are quantified on both the axonal field (AF, purple) and the dendritic field (DF, yellow); spread are estimated from the normalized SNR (right). **e**, Distributions of the positive peak amplitudes (top), the negative peak amplitudes (middle), and the spatial spreads (bottom). *** *p* = 9.3 × 10^−27^, 2.7 × 10^−31^, 0.641/6.2 × 10^−8^,5.8 × 10^−24^, 1.4 × 10^−24^/1.7 × 10^−34^, 7.8 × 10^−67^, 1.18 × 10^−22^, *n* = 134 TCAs, *n* = 715 V1Ns, *n* = 8 mice. **f**, Schematic of V1 tangential insertion from the side for pharmacological control experiments. **g**, A coronal slice showing the Neuropixels probe track (colored in yellow), muscimol injection in V1 (green), and TTX injection in dLGN (bottom, colored in magenta). **h**, Representative TCA (top) and V1N (bottom) with the corresponding MCW (left) and waveform at the best channel (inset). TCA and V1N activity over the time course of different pharmacological treatments (right), each shown at its best channels position (y-axis): control (gray), muscimol in V1 (green) and TTX in the dLGN (magenta). **i**, Corresponding firing rates of TCA and V1N during the different pharmacological conditions. *** *p*= 8.57 × 10^−16^,9.19 × 10^−16^/ 2.25 × 10^−51^,0.786, *n* = 86 TCAs, *n* = 301 V1Ns, *n* = 5 mice. **j**, Raster plot of TCA (top) and V1N (middle) firing to 100 trials. Comparison of Fano-Factor values in TCA and V1N. V1N shows higher variability in firing. *** *p* = 5.82 × 10^−10^, *n* = 134 TCAs, 715 V1Ns, *n* = 8 mice. Two-sided Wilcoxon signed-rank test for i, Two-sided Wilcoxon rank-sum test for e and j.

9.8 µV, DF = 12.12 ± 6.8 µV, *p values*: V1N to AF *p =* 9.3 × 10^−27^, V1N to DF *p =* 2.7 × 10^−31^, AF to DF *p =* 0.641, negative peak amplitude: V1N = -54.9 ± 35 µV, AF = -26.1 ± 13.9 µV, DF = -12.5 ± 7.9 µV, *p values*: V1N to AF *p =* 6.2 × 10^−8^, V1N to DF *p =* 5.8 × 10^−24^, AF to DF *p =* 1.4 × 10^−24^, two-sided Wilcoxon rank-sum test, *n* = 715 V1N, *n =* 134 TCAs, *n =* 8 mice). In addition, both AF and DF exhibit a wider spatial spread^15^ compared to V1N (Fig. 1e, spread: V1N = 299 ± 10.5 µm, AF = 303 ± 14.2 µm, DF = 448 ± 17.3 µm, *p values*: V1N to AF *p =* 6.2 × 10^−8^, V1N to DF *p =* 5.8 × 10^−24^, AF to DF *p =* 1.4 × 10^−24^, two-sided Wilcoxon rank-sum test, *n* = 715 V1N, *n =* 134 TCAs, *n =* 8 mice). Overall, the small amplitude of TCA waveforms combined with their larger spread suggests that high density of recording sites with low noise level^12^ are a prerequisite to detect TCA waveforms in the cortex *in vivo*.

To confirm that the TCA waveforms (Fig. 1d, left) reflect the extracellular electrical activity of thalamic axons in cortex, we performed several control experiments. This set of experiments shows that: 1-TCA waveforms do not stem from cortical neurons as upon muscimol injection (a GABA_A_ agonist) the TCA waveforms remain active (Fig. 1h-i, green, TCA firing rate: control = 16.9 ± 11.4 spikes/s, muscimol 3.49 ± 4.06 spikes/s, *p =* 8.57 × 10^−16^, two-sided Wilcoxon signed-rank test, *n =* 86 TCA, *n =* 5 mice) while the activity of V1N is blocked (Fig. 1h-i, green, V1N firing rate: control = 8.7 ± 10.1 spikes/s, muscimol 0.15 ± 0.7 spikes/s, *p =* 2.25 × 10^−51^, two-sided Wilcoxon signed-rank test, *n =* 301 neurons, *n =* 5 mice). Note that while TCA remain active, their firing rate is reduced upon muscimol injection, which could be due to the reduction of corticothalamic feedback^16,17^ or the activation of pre-synaptic GABA_A_ receptors in thalamic axons^18^. 2-TCA waveforms originate from the dorsal lateral geniculate nucleus (dLGN), as their activity is abolished upon injection of Tetrodotoxin (TTX, a blocker of sodium voltage-gated channels) in the dLGN (Fig. 1h-i, pink, TCA firing rate: muscimol = 3.49 ± 4.06 spikes/s, TTX = 0.019 ± 0.06 spikes/s, *p =* 9.19 × 10^−16^, two-sided Wilcoxon signed-rank test, *n =* 86 TCAs, *n =* 5 mice). 3-Finally, we quantified the trial-by-trial variability of the visually evoked activity and found higher variability in V1N represented by an increased Fano Factor compared to thalamic neurons, which reproduced previous results^19^ (Fig. 1j, Fano Factor: V1N = 1.12 ± 0.04, TCA = 0.71 ± 0.03, *p =* 5.8 × 10^−10^, two-sided Wilcoxon rank-sum test, *n* = 715 V1N, *n =* 134 TCAs, *n =* 8 mice).

Recording simultaneously the activity of thalamic axons and nearby V1 neurons permits to capture a large number of monosynaptically connected neuron pairs within individual recordings (Fig. 2d). Monosynaptic connections were identified by the presence of a transient and significant peak in their spike-train cross-correlograms (CCG, Fig. 2c/e, cf. methods). As expected, connected TCA-V1N pairs lie within close proximity in the cortical tissue (Fig. 2f, median distance within cortex = 62.1 µm, first quartile = 40 µm, third quartile = 121 µm, *n =* 208 connected pairs, *n =* 8 mice). This proximity of TCA to connected V1N explains why classical paired recordings require precisely aligned electrodes. The dense sampling of thalamic axons and cortical neurons by the high-density electrode resolves the challenging alignment step and thereby allows to efficiently map the thalamocortical connectivity on a large scale *in vivo* (mean number of identified connections per recordings: awake mice = 76.5 ± 30.7 connections, *n* = 2 mice; anesthetized mice = 9.3 ± 3.5 connections, *n* = 6 mice). Moreover, due to the high yield of identified connections, this approach allows to unravel divergent (mean identified diverging contacts between single TCA and V1N = 3.41 ± 0.05 connections) and convergent connections (mean identified converging contacts between TCA and single V1N = 3.1 ± 0.03 connections) (Fig. 2g). Finally, we can estimate the spike transmission efficacy^13^ providing an *in vivo* measure of the thalamocortical connection strength (Fig. 2h, median efficacy = 1.8 %, first quartile = 1.2 %, third quartile = 3.1 %, n = 208 connected pairs, n = 8 mice).

**Figure 2:**
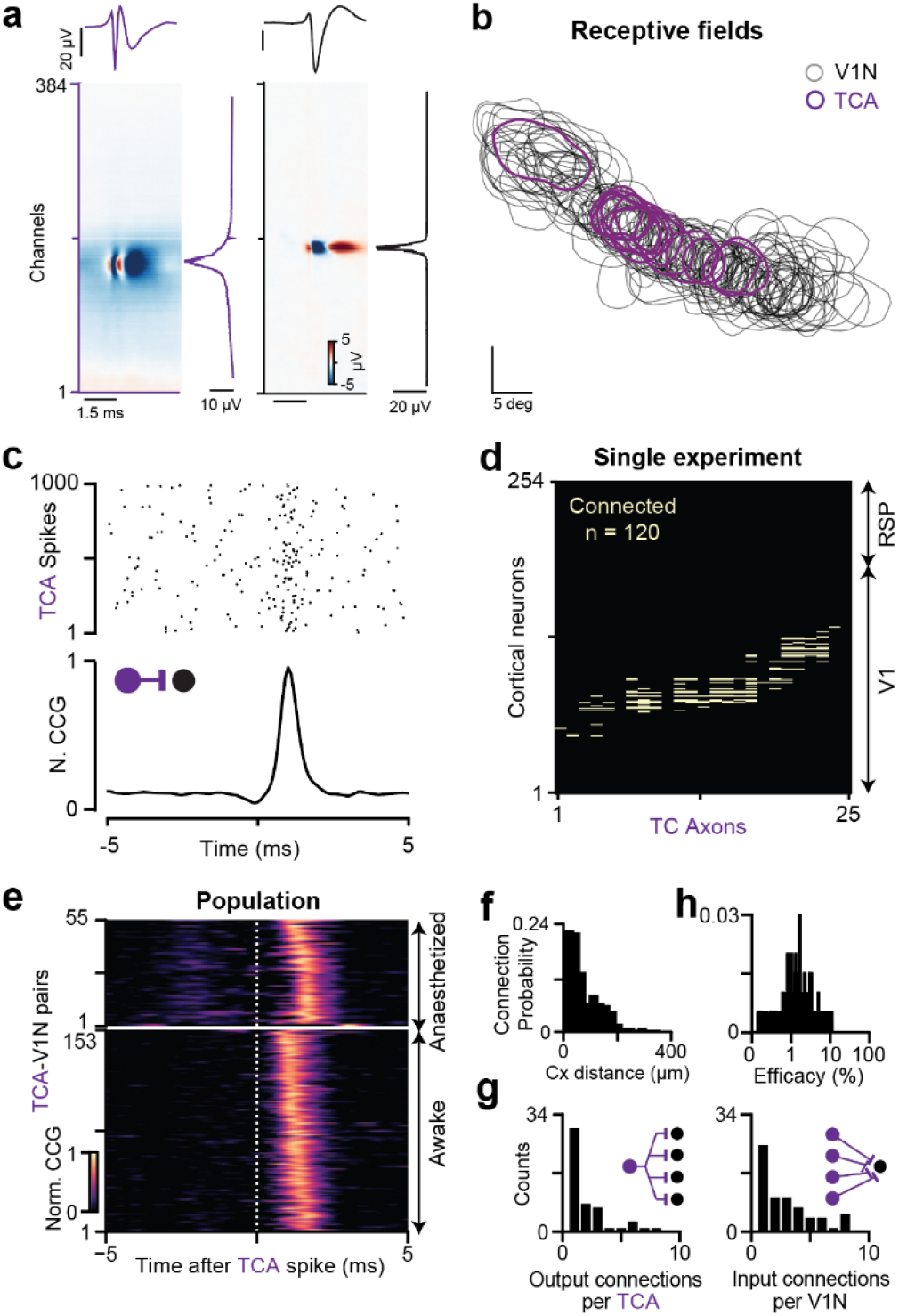
Efficient mapping of thalamocortical monosynaptic connectivity in anesthetized and awake mice. **a**, Multi-Channel-Waveform (MCW) of a TCA (left) and a neighboring V1N (right) with their respective amplitude profiles (righter insets). **b**, Overlapping receptive fields of a population of TCA and neighboring V1N. **c**, Raster plot showing V1N responses of a connected TCA-V1N pair triggered on 250 independent TCA spikes (top) and the corresponding CCG (bottom). **d**, Connectivity matrix from a single awake recording containing 120 simultaneously detected monosynaptic contacts between 25 TCAs and 211 V1Ns. **e**, CCG profiles of all measured monosynaptic connections in anesthetized (*n* = 56 connections, *n* = 6 mice) and in awake mice (*n* = 153 connections, *n* = 2 mice). **f**, Distribution of peak channel distances between the connected TCA and V1N. **g**, Distributions of the diverging TCA (left) and converging V1N (right) connections. **h**, Distribution of connection efficacy on log axis.

In this work, we demonstrate that tangential insertions of high-density electrodes into cortical layer 4 of awake or anesthetized head-fixed mice is a reliable and efficient method for mapping the thalamocortical connectivity *in vivo*. Thus, placing high-density electrodes tangentially into cortex is a promising approach for revealing multiple thalamic inputs to cortical neurons in possibly most contexts of rodent behaviors.

## MATERIAL AND METHODS

### Animals, surgery, insertion, and staining

#### Legal bindings

Agreements from the local authorities were obtained for the anesthetized (*n =* 6) and the awake head-fixed (*n =* 2) experiments (Landesamt für Gesundheit und Soziales Berlin - G 0142/18). Maximum attention was focused to minimize the number of used animals. Adult C57BL/6J mice were obtained from local breeding facility (Charité-Forschungseinrichtung für Experimentelle Medizin). <u>For the two awake mice</u>, a head-post was implanted 1.5 weeks prior to the recording under isoflurane anesthesia, painkiller was provided for 3 days after implantation, then mice were habituated with daily sessions gradually increasing duration on the set-up including exposure to visual stimuli.

<u>On the recording day</u>, the mouse was induced with controlled level of isoflurane (2.5% in oxygen Cp-Pharma G227L19A). The mouse underwent surgery in a stereotactic frame (Narishige) where the isoflurane level was gradually lowered (0.7-1.5%) with a constant monitoring of the absence of both responses to tactile stimulations and vibrissa twitching. <u>During surgery,</u> the temperature was constantly monitored (FHC-DC), the eyes were covered with ointment (Vidisic), meanwhile a head-post was positioned, and a dental cement-based crown (Paladur, Kuzler) was built around the skull including a grounding wire. For the tangential insertion from the side, enough room was prepared on the lateral side of the crown for the insertion to enter the brain below the lateral cranial ridge. Once the head-post was solidified, and all craniotomies were finished for the desired experiment(s), the mouse was transferred into the set-up to proceed with the recording. For the awake recordings, the mouse recovered from craniotomy surgery for one to two hours before proceeding to the experimental set-up.

<u>Two different insertions</u> are proposed to reliably achieve TCA measurement in V1: a) the insertions from the top (Fig. 1a, Fig. 2a-e) - advised for awake recordings - with a negative 25-30° from the azimuthal plane (within a coronal plane) or b) the insertions from the side - advised for anesthetized pharmacological controls - (Fig. 1a) with a positive 15-20° from the azimuthal plane (within a coronal plane). With such titled insertion(s), to avoid any slight bending of the probe upon tissue contact, removing temporarily the grounding solution will be required. <u>All stereotactic coordinates are reported from lambda,</u> in the antero-posterior (AP), the medio-lateral (ML), and the dorso-ventral (DV) axis. Insertions from the top enter the brain at 0 to 900 μm AP, +250 to 0 μm ML and -100 to -500 μm DV. While insertions from the side enter the brain at 0 to 900 μm AP, -5000 to -4000 μm ML and -2000 to - 500 μm DV from lambda. In both implantations, the probe was inserted for at least 4000 µm, then withdrawn by 50 µm to release mechanical pressure. After the probe has settled in the tissue for ∼10-20 minutes, a short sparse noise stimulus was displayed to produce online RF mapping. This first short recording was also used to probe the placement of the tangential insertion within the cortical layer 4 (cf. below semi-online detection of TCA). If both RF mapping and TCA waveform identification (Extended data 1) appeared in similar channels, <u>the data acquisition was started</u>. A set of visual stimuli set was presented, followed either by the different pharmacological applications (cf. below) or by repeating the set of visual stimuli in the awake mice. Once the recording is terminated, the probe was partially removed and re-inserted, coated with DiI (Abcam-ab145311, in EtOH). <u>The mouse was then sacrificed</u> with an excess of isoflurane (>4%) followed by a cardiac perfusion with phosphate buffer saline solution (PBS) and 4% paraformaldehyde (PFA). The brains were left in PFA overnight and stored in PBS until slicing (Leica VT1200 S) and mounting (100 µm slices, DAPI-Fluoromount-G Biozol Cat. 0100-20). The obtained mounted brains were further analyzed <u>to perform the 3D reconstruction</u> of the probe’s exact tissue location. To do so, we used SHARP-track^20^ in order to relate our staining to the Allen Mouse Brain Common Coordinate Framework. We used the entry and exit point of visually driven activity from the MUA as the outer boundaries of the visual cortex (Fig. 1b in red) to align and scale to the cortical layers and regions identified by SHARP-track. Consequently, each channel on the probe was assigned to a tissue location, to determine the location for each V1N and TCA (Fig. 1c).

### Pharmacological applications

In the majority of the dataset (5 out of 8 recordings), <u>pharmacological solutions</u> were applied to confirm the origin of the measured TCA waveforms in V1. A first set of visual stimuli set was presented before the injector (Drumond, Nanoject II) was lowered vertically right next to the probe at 500 to 1000 µm ML, 400 to 0 μm AP, -1100 to -1400 μm DV from lambda, to inject 100 nL of muscimol solution (Abcam, ab120094, 2.5 mM in PBS) containing a dye (Cholera Toxin subunit B, Alexa 488 Conjugate -C22841, Invitrogen). Upon muscimol injection and diffusion (∼5 minutes), a clear reduction of ongoing spontaneous cortical activity should be observed. The same visual stimulus set was re-exposed followed by a second pharmacological injection of TTX (100 nL, Biozol-HB1034, 100 μM in PBS) with the same dye in the dLGN at 2500 µm ML, 2000 μm AP, -4000 μm DV from lambda. A last repeat of the chirp stimulus was presented after injection of TTX to confirm the loss of visually driven activity in V1. As the same staining was used in both injection, pseudo colors were arbitrarily attributed in the different brain regions for easier illustrations (Fig. 1g).

### Hardware, software and visual stimulation

<u>Neuropixels probes 1 signal was acquired</u> on a PXIe system (National Instrument NI-PXIe-1071) using Open Ephys acquisitions software (www.open-ephys.org). The raw signal was split and stored in the local field potential (0-300 Hz) and the action potential (AP) band (300 Hz to 3 kHz). Ecephys was used to calculate standard cluster quality metric (Extended Data Fig. 3, https://github.com/AllenInstitute/ecephys_spike_sorting). Except for sorting with Kilosort^21^ (https://github.com/MouseLand/Kilosort) which was done in MATLAB 2018 & 2019 (www.mathworks.com), <u>all data analysis was performed in Python 3</u> (www.anaconda.com); statistical tests were performed using either the two-sided Wilcoxon rank-sum test except for the pharmacological recording where we use a two-sided Wilcoxon signed-rank test (Fig. 1i). P values are indicated as following: “***” indicates a *p value* below 0.001. <u>The visual stimuli</u> were displayed on a monitor screen (Dell, calibrated at refresh rate = 120 Hz, mean luminance = 120 cd/m^2^). The stimulus set was made of: two sparse noise stimuli with either dark or light targets (15 deg size, 2 targets/frames of 100 ms, 20 repeats for each position on a grid of 36×22 squares), moving bars (white bars on black background, 10 deg in width, 12 directions, fixed speed of 90 deg/s), followed by the final full-field chirp stimulus^13^. For each exposed image, a logic electric signal was produced and sent to the digital-to-analog converter (NI-DAQmx connected in the NI-PXI with a PXI-8381) to support later re-alignment and re-interpolation of the time between the visual stimuli and the registered spike times.

### Semi-online detection of TCA

To ensure the tangential insertion of the Neuropixels probe through the cortical layers 4, and the quantity of TCA in the recording, we strongly advise to look at RF mapping and TCA waveforms plotting together. To do so, the short sparse noise stimulus was displayed and the neuronal activity recorded^22^. <u>The first main verification is to use an adaptation of the “make_fig.m” script from Kilosort</u> (Extended Data Fig. 1, compatible with KS 2 & 2.5) (https://github.com/KremkowLab/Cortical-Axon-on-Neuropixels-in-Kilosort) to directly examine in response to the short sparse noise stimulus the possible presence of TCA waveforms^23^. This script allows to visualize waveforms with the highest rebounds as the most likely clusters being axons (Extended Data Fig. 1a) to illustrates their locations on the probe (Extended Data Fig. 1c, in red). Due to cortical mantle curvature; it can be quite tricky to achieve an extended layer 4 tangential insertion capturing large populations of TCAs and V1Ns. Obtaining numerous axons (n > 50) during this verification step is therefore strongly advised (Extended Data Fig. 1b); re-insertion can often be needed to achieve a higher yield. Trustable axons usually have a rebound of sizes reaching -0.2 to -0.3 in the amplitude axis (Extended Data Fig. 1c purple area). <u>Second</u>, a more classical online analysis <u>will confirm the placement of the probe in the visually driven channels</u> by mapping receptive fields (RF) as published recently by the lab^22^.

Altogether, the combination of RF mapping, indicating visually driven activity on a given set of channels, together with the Kilosort plotting, indicating possible TCA waveforms <u>on the same visually driven channels</u>, will both together give the confirmation that the tangential insertion is properly placed next to the layer 4 of visual cortex, capturing visually driven TCA. For other sensory modalities adapted stimuli should be employed to confirm the overlapping sensory responses to the location of the measured TCA.

### Spike sorting, axonal detection, and characterization

<u>Kilosort 2.5</u> was used for spike sorting. Manual curation was done post-hoc using Phy2 (https://github.com/cortex-lab/phy/). During curation, it is advised to avoid rejection based on multiple spatial peaks and too large spread^24^; furthermore, given the smaller amplitudes of TCA waveforms (Fig. 1e), it is advised to ignore the “MUA” labelling in Phy2 when considering a putative TCA cluster. Once the manual curation is finished, quality control should be applied such as removal of double-counted spikes, inter-spike-interval violations (> 0.05%) and isolation distance (isolation distance > 10 a.u.), Extended Data Fig. 3)^13^. Quality metrics were obtained from ecephys (https://github.com/AllenInstitute/ecephys_spike_sorting). <u>We evaluated the multi-channel waveforms</u> (MCW) by averaging the raw action potential signal on all spike times from each individual single unit (up to 50 000 spikes), followed by a channel time offset correction of 2.78 μs between neighboring channels of the same analog-digital-converter^13^. Consequently, the MCW represents the spatiotemporal profile of the electrophysiological signal associated to the given single-unit, over the entire tissue covered by the Neuropixels probe (Fig. 1d, Fig. 2a, Extended Data Fig. 2c). <u>In the pharmacological experiments</u> (*n =* 5 mice), an extra stability control is strongly advised to reject bad clusters including waveforms of different shapes in the different pharmacological conditions. This step was performed in a custom written GUI illustrating the MCW recalculated in each pharmacological condition separately^13^. Once all controls are finished, <u>the amplitude and the spread of each cluster can be estimated</u>. The Signal-over-Noise-Ratio (SNR) is calculated as the variability of the maximal amplitude in each channel. This SNR over all channels is then normalized to quantify the spread size as the number of channels above 0.1 (Fig. 1d-e). When the probe is by chance aligned with the TCA axonal path running below cortical layer 6, the action potential propagation speed can be estimated (Extended Data Fig. 2)^13^. <u>Axonal propagation speed was measured</u> using a minimal window of 5 channels away from the best channel and 5 channels away from the Neuropixels tip. Within this window, the action potential peak was detected in a fixed time windows (−2.5 to -0.1 ms) just when its amplitude reached 4*std of the measured baseline (−5 to -2.5 ms). A line was fitted between the different action potential peak times in neighboring channels to determine the propagation speed; interpolation values above R = 0.725 were kept for quantification. This step was repeated on each of the four Neuropixels electrode columns independently; the best fit value was kept obtaining 7 distinguishable axonal paths showing an average of 0.92 ± 0.6 m/s propagation speed. It is also possible to distinguish <u>different firing properties of spiking between TCA and V1N</u> by looking into their spiking variability. The variability of spiking is quantified by the different responses to a 1 s alternating 10° checker stimulus. The responses of TCA and V1N to 100 repeated trials (Fig. 1i top and middle) illustrates the higher trial-to-trial variability of V1N. This inter-trial variability is divided by the average firing during the same period to estimate the Fano Factor of each V1N or TCA^19^.

### Receptive fields and functional synaptic connectivity

To analyze visual responses properties, the RFs were first calculated from the PSTH produced by summing the activity on each exposed pixel over the entire stimulation. Once this PSTH is produced, the RFs are the summation over time of the firing evoked on each pixel by the visual stimulus^7^. <u>RF contours were interpolated</u> by a factor of two using the 2D-cubic-interpolation function from SciPy package; only clusters with RFs showing a SNR (1/std (RF)>15) were kept for quantification (Fig. 2c, Extended Data Fig. 3a).

Monosynaptic connections between TCA and V1N was detected using established methods^13^. Cross correlations were calculated using pycorrelate (https://github.com/tritemio/pycorrelate). To estimate the jitter, spike times over the entire recording were randomized within a consecutive 10 ms window and subtracted from the CCG. <u>Reliable connections were detected</u> from significant peaks within a window (0.5 to 4 ms for at least 4 consecutive time bins) above the baseline (3 x std, -4.5 to 0.5 ms, Fig. 2c). The distance on the probe between the TCA and the V1N was calculated between their respective peak channels (Fig. 2f). Their synaptic efficacy was estimated from the jitter-corrected CCG as the area under the monosynaptic peak divided by the total number of spikes from the TCA^13^.

## Data availability

A preliminary recording has been made available online^23^.

## Code availability

The analysis used for the visualization of putative TCAs is available on the laboratory GitHub repository^23^. Furthermore, the online analysis used to represent the visually driven MUA activity has been published in a separate publication^22^.

## Acknowledgements

We thank J. Poulet, B. Judkewitz for helpful discussions and materials exchanges during the project; Perrenoud Q., Lupashina T. and Kosubek-Langer J. for comments on the manuscript. The mouse head schema used in Fig. 1f was adapted from https://doi.org/10.5281/zenodo.3925903. This work was supported by the DFG Emmy-Noether grants KR 4062/4–1 and KR 4062/4–2 (JK).

## Authors contribution

J.S. and J.K. conceived and designed the study; J.S., C.G., collected the data; J.S., and J.K. analyzed the data and J.S. and J.K. wrote the manuscript with inputs from all authors.

## SUPPLEMENTARY MATERIAL

**Extended Data Fig. 1:**
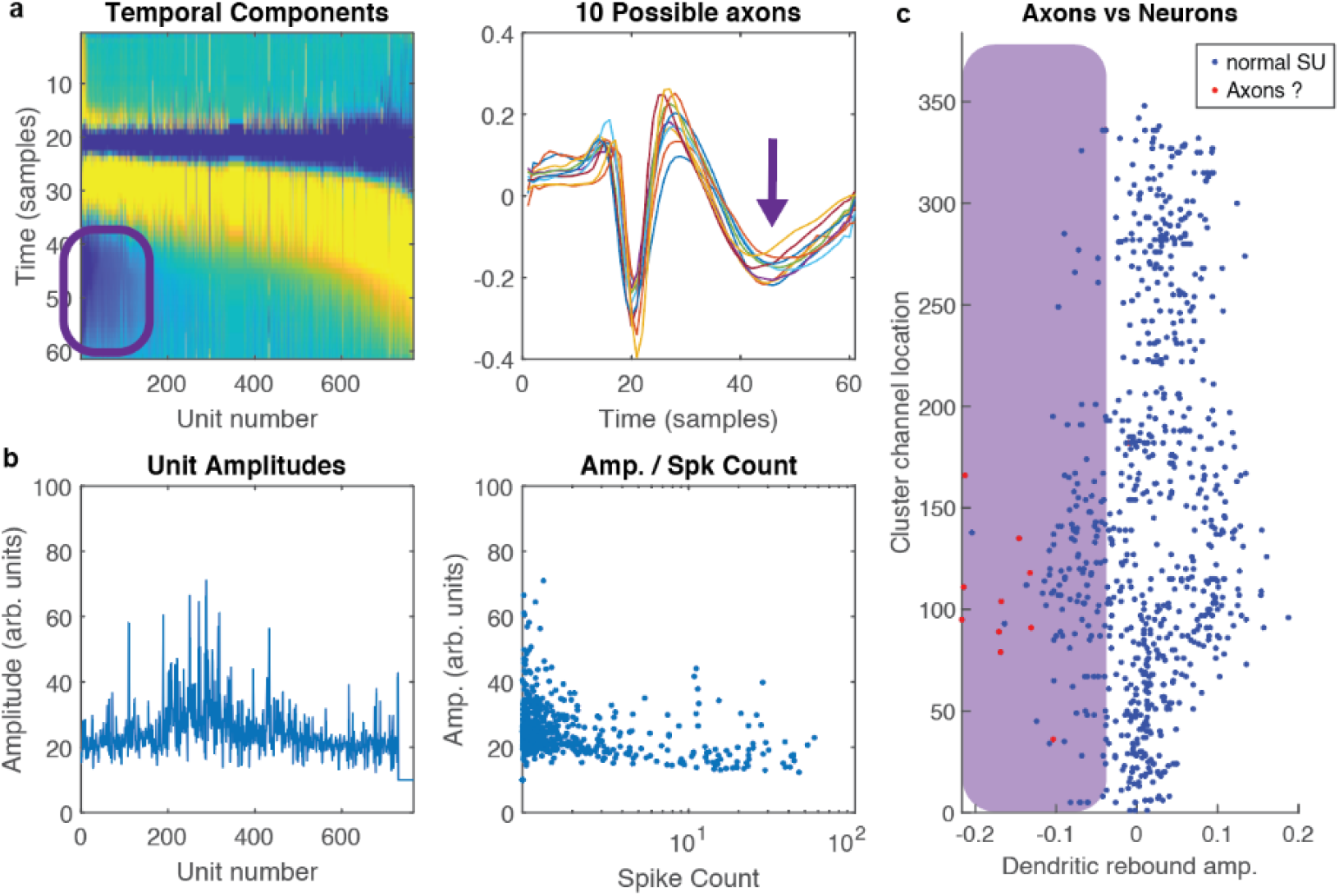
Adaptation of Kilosort to estimate the presence of TCA in a recording. **a**, Modified plots from the temporal components showing the captured waveforms, sorted by the amplitude of their rebound. Putative axons are located on the left with their negative rebound values highlighted in purple. Visualization of the first 10 waveforms with the highest rebound (right). **b**, Original Kilosort plots showing amplitude and spike count of the captured units. **c**, The channel locations of each unit is shown on the probe against their rebound amplitude in order to visualize the locations of putative axons (purple area). Note: the 10 clusters with highest rebound amplitude (from **a**) are highlighted in red.

**Extended Data Fig. 2:**
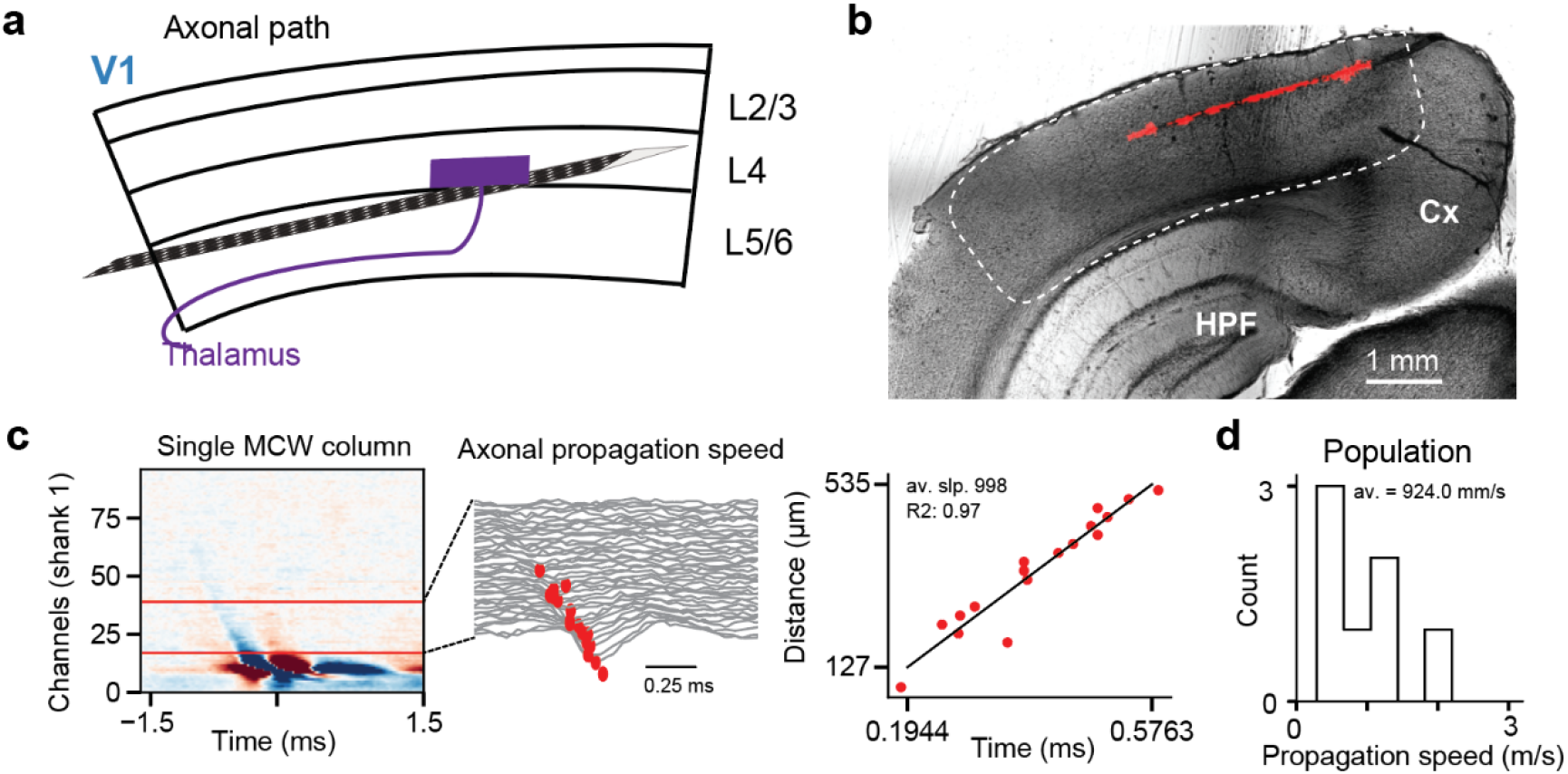
TCA propagation speed. **a**, Schematic of an insertion from the side. **b**, Coronal slice with Neuropixels probe track (in red). **c**, Quantification of the axonal propagation speed from the MCW (left). Zoom of the measured time points on which the axonal “tail” can be reliably detected (middle); together with the corresponding interpolation (right). **d**, Distribution of the conduction velocity (1 m/s, *n =* 7 TCAs, *n =* 3 mice).

**Extended Data Fig. 3:**
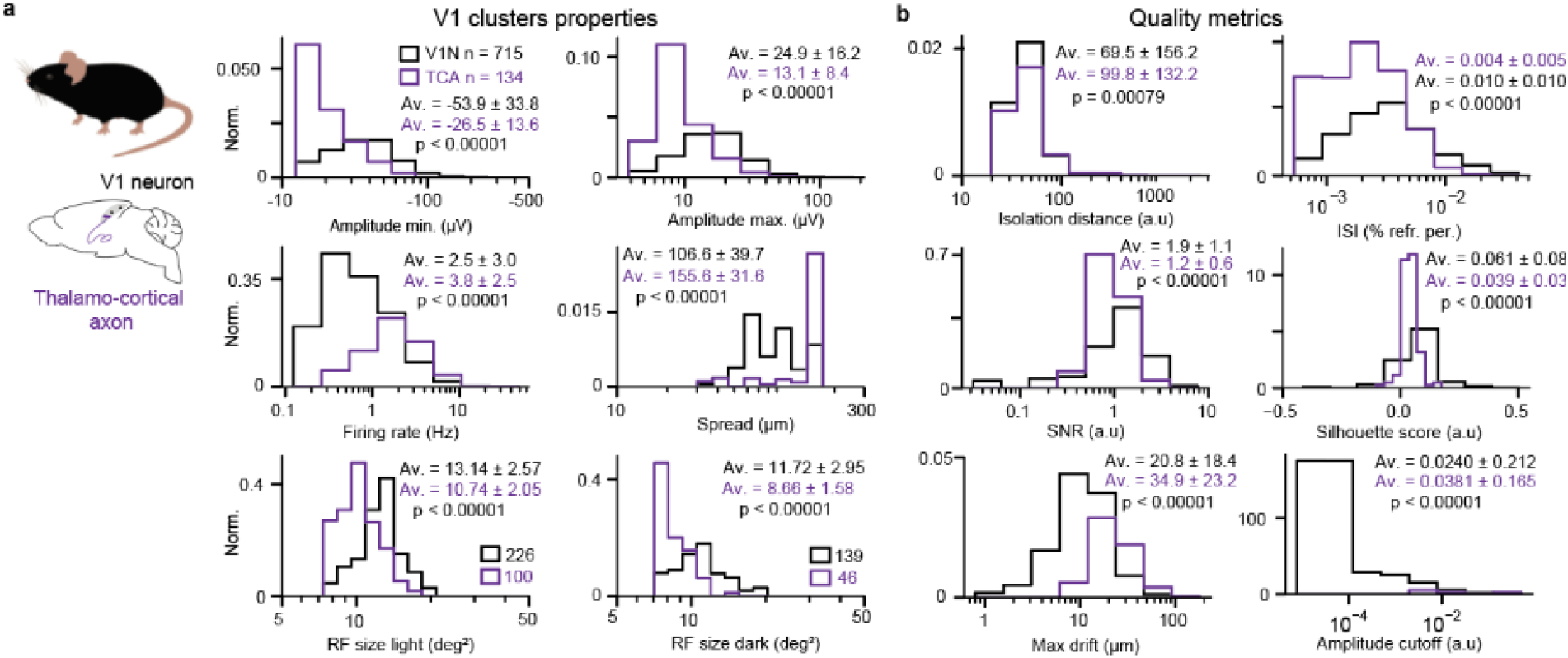
Properties and quality metrics of TCA & V1N. **a**, Cluster properties of V1N (black) and TCA (purple): Amplitudes are obtained from ecephys (negative - top left, positive - top right), both on log axis. Spread (middle right) and firing rate (middle left). Receptive field (RF) sizes were estimated from high SNR clusters (SNR > 15). **b**, Quality metrics for Corresponding TCA and V1N.

